# PWWP INTERACTOR OF POLYCOMBS (PWO1) links PcG-mediated gene repression to the nuclear lamina in *Arabidopsis*

**DOI:** 10.1101/220541

**Authors:** Pawel Mikulski, Mareike L. Hohenstatt, Sara Farrona, Cezary Smaczniak, Kerstin Kaufmann, Gerco Angenent, Daniel Schubert

## Abstract

Polycomb group (PcG) proteins facilitate chromatin-mediated gene repression through the modification of histone tails in a wide range of eukaryotes, including plants and animals. One of the PcG protein complexes, Polycomb Repressive Complex 2 (PRC2), promotes repressive chromatin formation via tri-methylation of lysine-27 on histone H3 (H3K27me3). The animal PRC2 is implicated in impacting subnuclear distribution of chromatin as its complex components and H3K27me3 are functionally connected with the nuclear lamina (NL) - a peripheral protein mesh that resides underneath the inner nuclear membrane (INM) and consists of lamins and lamina-associated proteins. In contrast to animals, NL in plants has an atypical structure and its association with PRC2-mediated gene repression is largely unknown. Here, we present a connection between lamin-like protein, CROWDED NUCLEI 1 (CRWN1), and a novel PRC2-associated component, PWWP INTERACTOR OF POLYCOMBS 1 (PWO1), in *Arabidopsis thaliana*. We show that PWO1 and CRWN1 proteins associate physically with each other, act in the same pathway to maintain nuclear morphology and control expression of similar set of target genes. Moreover, we demonstrate that PWO1 proteins form speckle-like foci located partially at the subnuclear periphery in *Nicotiana benthamiana* and *Arabidopsis thaliana*. Ultimately, as CRWN1 and PWO1 are plant-specific, our results argue that plants developed an equivalent, rather than homologous, mechanism of linking PRC2-mediated chromatin repression and nuclear lamina.

## Introduction

Polycomb group (PcG) proteins facilitate chromatin-mediated gene repression by modifying histone tails in a wide range of eukaryotes, including plants and animals. A subset of PcG proteins constitutes Polycomb Repressive Complex 2 (PRC2). A canonical form of PRC2 was initially discovered in *Drosophila melanogaster*, where it consists of: E(z) – a component with catalytic functions, Esc –a WD40 motif-containing scaffolding protein, Su(z)12 – a Zinc-finger subunit facilitating binding to nucleosomes, and p55 – a nucleosome remodelling factor (Schwartz and Pirrotta, 2007). PRC2 is present also in the model higher plant, *Arabidopsis thaliana*, albeit with several components being multiplicated and partially diversified in expression pattern and set of targets. PRC2 in *Arabidopsis thaliana* is made up of: CLF/MEA/SWN - E(z) homologs, VRN2/EMF2/FIS - Su(z)12 homologs, FIE - ESC homologs, and MSI1-5 - p55 homologs (Hennig and Derkacheva, 2009). PRC2 mediates its repressive function by catalysing methylation of lysine 27 on histone H3 (H3K27), which is in turn recognized by reader proteins that silence underlying DNA sequences (Xiao et al., 2016). Depending on the species, PRC2 methylates H3K27 in various contexts (Ebert et al., 2004; Ferrari et al., 2014; Jacob and Michaels, 2009), with H3K27 trimethylation (H3K27me3) being the most typical modification.

PRC2 functions on different layers of gene repression (reviewed in (Del Prete et al., 2015)), including regulation of target spatial distribution in the nucleus, namely at the nuclear periphery. Nuclear periphery is a subnuclear space in the vicinity to nuclear envelope (NE), a double lipid bilayer forming inner and outer nuclear membrane (INM and ONM, respectively), and nuclear lamina (NL), a nuclear protein mesh physically associated with INM. NL comprises of lamin-associated membrane proteins and lamins themselves, the latter one being categorized as A- and B-type, depending on the structural properties and expression pattern (Dechat et al., 2010). Apart from a role in the nuclear architecture, both lamin types were implicated to influence subnuclear organization of the chromatin, exemplified by the existence of lamin-bound chromatin regions, called lamina-associated domains (LADs) in animals.

Several lines of evidence indicate that PRC2 plays an important role in the regulation of LADs and peripheral localization of specific chromatin types. These include: (1) H3K27me3 enrichment at tissue-specific LADs (variable LADs, vLADs) and the borders of constitutive LADs in mammalian cell cultures (Fakhouri et al., 2010; Harr et al., 2015); (2) H3K27me3 enrichment at chromatin domains bound by LEM-2, lamina-associated protein, in *C. elegans* (Ikegami et al., 2010); (3) H3K27me3 accumulation at transgene multicopy arrays that are associated with NE in *C. elegans* (Meister et al., 2010; Towbin et al., 2010); (4) decrease of peripheral localization for vLAD fragment upon knockdown of EZH2, a mammalian homolog of E(z), in mouse fibroblast (Harr et al., 2015) and (5) recruitment of vLAD fragment to NL mediated by Ying Yang 1 (YY1), an interactor of PRC2 (Harr et al., 2015).

However, general H3K27me3 occupancy and PRC2 targeting are not the exclusive determinants of chromatin association with animal lamina. On one side, a number of PRC2-independent pathways was reported to be also involved in that process (Harr et al., 2016), with some being specific to cell type or the species. On the other, only a subset of all PRC2 targets can be found in LADs (Hänzelmann et al., 2015) and spatial distribution of PRC2 components or H3K27me3 in some tissue types concerns localization to both, nuclear periphery and nuclear interior (Eberhart et al., 2013; Luo et al., 2009; Wu et al., 2013). Therefore, understanding the role of PRC2 in peripheral positioning requires characterization of target-specific PRC2-associated proteins and deciphering the interplay with other factors involved in tethering to NL.

In plants, general impact of chromatin on gene spatial distribution and contribution of Polycomb group-mediated repression in this process was investigated only in a handful of studies (Pecinka et al., 2002; Rosa et al., 2013; Rosin et al., 2008) and remains largely unknown. In fact, no lamina-associated domains were identified in the green lineage and it was long believed that plants do not have the lamina, as they lack sequence homologs of the lamin proteins (Melcer et al., 2007). Instead, plant NL contains the lamina-like proteins (Ciska and Moreno Díaz de la Espina, 2014), which, similarly to their animal counterparts, possess coiled-coil protein motif and show general localization to the nuclear periphery (Ciska and Moreno Diaz de la Espina, 2013; Gardiner et al., 2011). Lamina-like genes in *Arabidopsis* form plant-specific family CROWDED NUCLEI (CRWN), containing 4 members (*CRWN1-4*). The most prominent phenotypes of CRWN family loss-of-function mutants are: reduced nuclear size, increased nuclear DNA density and abnormal nuclear shape, with the most pronounced effect seen in *crwn 1 crwn2* (Dittmer et al., 2007; Sakamoto and Takagi, 2013; Wang et al., 2013). Furthermore, CRWN proteins associate with a range of other plant NE/NL components (Goto et al., 2014; Graumann, 2014). Interestingly, CRWN genes also affect constitutive heterochromartin organization and 3D chromosome arrangement - different CRWN mutant combinations show altered chromocentres’ integrity (Dittmer et al., 2007; Wang et al., 2013) and *crwn1 crwn4* exhibits elevated *trans*-chromosomal interactions, but unchanged contact frequency *in cis* (Grob et al., 2014). However, the exact mechanism of CRWN proteins’ function in chromatin organization remains unknown.

In our previous work, we identified *Arabidopsis* protein PWWP INTERACTOR OF POLYCOMB1 (PWO1) as a novel plant-specific PRC2-associated factor. We showed that PWO1 interacts physically with PRC2 methyltransferases (MEA, SWN, CLF), displays epistatic interaction with CLF, controls expression of several PRC2-dependent target genes and is needed for full H3 and H3K27me3 occupancy at several PcG target genes (Hohenstatt, 2012).

Here, we present a connection between PWO1 and the plant nuclear lamina. We demonstrate that PWO1 interacts with CRWN1 physically and genetically, and that they both control expression of a similar set of genes. Furthermore, we show that PWO1 regulates nuclear size and partially localizes to peripheral speckles in the nucleus. Taken together, our results provide a putative link between PcG-mediated gene repression and chromatin organization at the subnuclear periphery.

## Results

### PWO1 physically associates with NL/NE proteins

After initial characterization of PWO1 (Hohenstatt, 2012), we sought to identify its protein interactors by unbiased quantitative proteomics. We undertook co-immunoprecipitation experiment coupled with mass spectrometry using a bait of PWO1-GFP fusion protein in *PWO1::PWO1-GFP Arabidopsis* transgenic line (Hohenstatt et al, 2017). The protein abundance was scored via label-free quantification (LFQ) analysis and a comparison to background sample (Col-0, wildtype) was used to identify significantly enriched proteins in the *PWO1::PWO1-GFP* line. As a result, we obtained a list of 109 putative PWO1 interactors. Interestingly, ∼ 60% of those overlap with putative components of crude plant nuclear lamina fraction (Sakamoto and Takagi, 2013) (Fig.1A), including constitutive NL/NE members and chromatin-associated proteins (Fig.1B, Table S1). Our attention was drawn to the presence of CROWDED NUCLEI 1 (CRWN1), a nuclear lamina protein with prominent role in nuclear morphology (Wang et al., 2013) that affects chromocenter organization (Dittmer et al., 2007) and interchromosomal contact frequencies (Grob et al., 2014).

**Fig.1.**
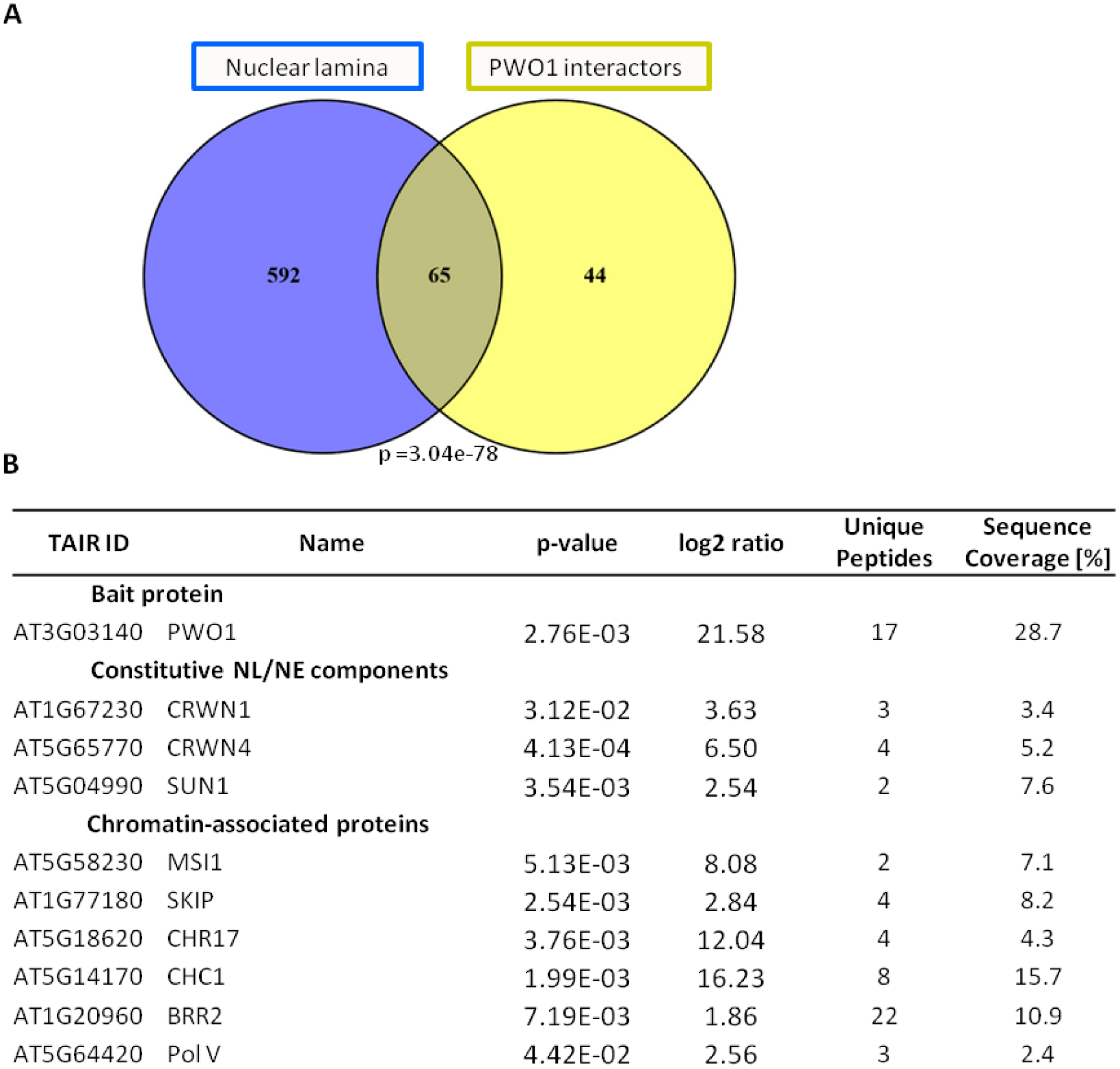
Identification of PWO1 interactors. (A) An overlap between *Arabidopsis* nuclear lamina (NL) fraction (Sakamoto and Takagi, 2013) and PWO1 interactors. PWO1 interactors correspond to proteins showing significant (p<0.05) enrichment over the wildtype in CoIP-MS/MS experiment with PWO1-GFP (*PWO1::PWO1-GFP* line) used as a bait. Significance level of the overlap was calculated using the Student’s t-test. (B) Mass spectrometry results for bait protein and selected candidates from an overlap of PWO1 interactors and NL components. For the full list of 65 overlapped proteins, see Table S1.

In order to confirm PWO1-CRWN1 interaction, we performed Yeast-Two-Hybrid (Y2H) and Acceptor Photobleaching FRET (FRET-APB) experiments. Both methods revealed no or very low interaction between CRWN1 and full-length PWO1, but much stronger association when only a C-terminal PWO1 fragment (634 – 769 aa) was used instead (Fig.2, Fig.3, Fig.S1). Such result might indicate steric constrains of overall PWO1 protein fold caused by introduction of fusion tags (Gal4-domains or FRET fluorophores) and masking of CRWN1-binding site in full-length PWO1 construct versions. In contrast, the usage of truncated PWO1 protein containing CRWN1-interaction region might resolve potential steric effects. We exclude the possibility of masking the interaction fragment by Gal4-domain or fluorophore tags as our Y2H and FRET assays were done on N‐ and C-terminal PWO1 fusion, respectively. The interaction with CRWN1 was shown to be specific to C-terminal part of PWO1 and not a simple outcome of random binding of any truncated PWO1 versions, as the other PWO1 fragments (frag.1 and 2) show still a background-level interaction (Fig.2B).

**Fig.2.**
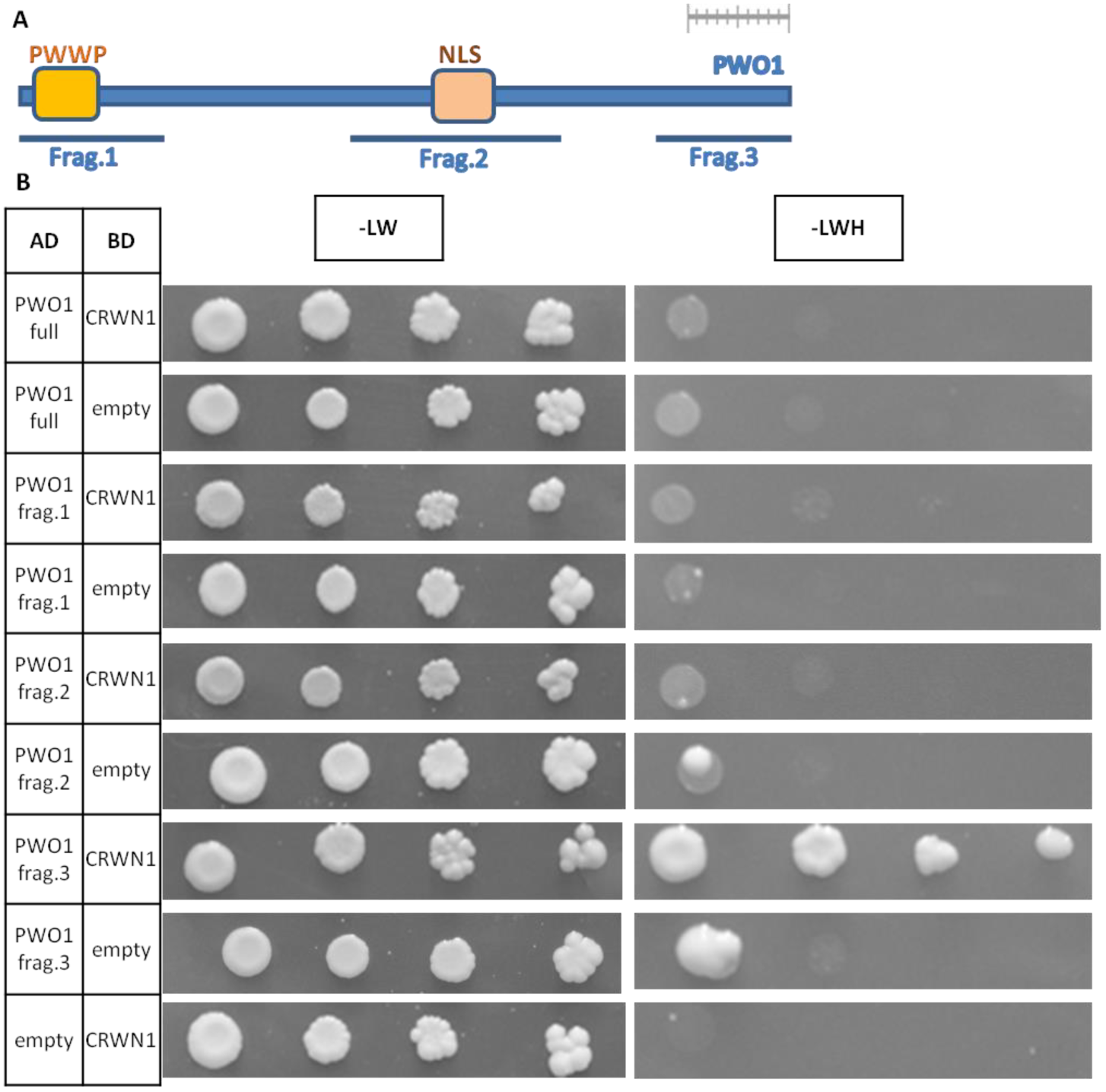
Yeast-two-hybrid (Y2H) analysis on PWO1-CRWN1 interaction. (A) Scheme of PWO1 protein sequence with marking of PWO1 fragments used for Y2H experiments. Scale bar on the top of the scheme corresponds to 100 aminoacids. (B) Y2H experiment. Y2H constructs containing inserts N-terminally fused to Gal4 activating domain (AD) or Gal4 binding domain (BD) were transformed in AH109 S.cerevisae strain. Coding sequences of full length CRWN1 (CRWN1), full length PWO1 (PWO1 full) and PWO1 fragments (PWO1 frag.1-3) were used as the inserts. Transformation with insert-containing constructs and empty vectors served as a negative control. Overall yeast growth and protein interaction between CRWN1, PWO1 and PWO1 fragments was assessed on transformation medium (-LW) and selection medium (-LWH).

**Fig.3.**
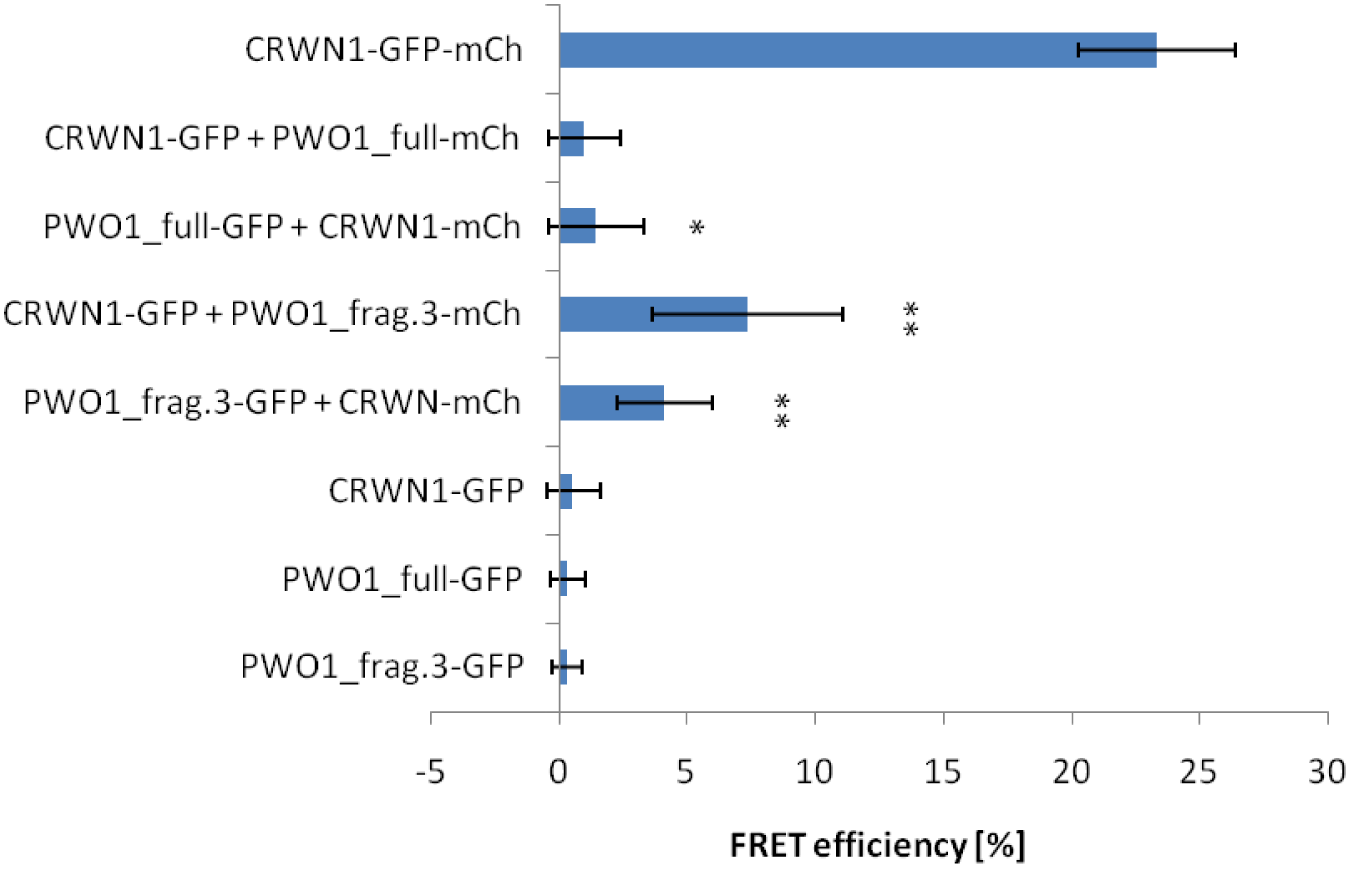
FRET-APB-based interaction between PWO1 and CRWN1. FRET efficiency was calculated for nuclei from the positive control (n=10), negative controls (n=15−17) and interaction samples (n=15−27). Error bars represent standard deviation values. Significance between interaction samples and their respective negative controls was inferred from Student’s t-test. Single asterisk (*) represents significance threshold: p<0.05, double (**): p<0.01.

Interestingly, the CRWN1-interaction region in PWO1 sequence does not contain any annotated protein domain (Fig.2A). However, an alignment to the closest PWO1 plant homologs revealed a high level of conservation for PWO1 C-terminal sequence in *Brassicaceae* species other than *Arabidopsis* (Fig.S2), highlighting a functional importance of this fragment. In contrast, PWO1 C-terminal fragment is largely absent from PWO2 and PWO3, suggesting partially separate functions between PWO family proteins in *Arabidopsis.*

### PWO1 interacts genetically with CRWN1

Given the physical association of PWO1 and CRWN1, we sought to investigate their functional connection by studying *pwo1*, *crwn1* and *pwo1 crwn1* mutants on the phenotypic level. As PWO1 is PRC2-associated protein (Hohenstatt, 2012), we included also *swn* and *swn crwn1* transgenic lines in our analysis to characterize the link between nuclear lamina and canonical PRC2.

A prominent feature of mutants in NE or NL components is the reduction in nuclear size and change of nuclear shape. In order to characterize alterations in nuclear morphology, we isolated whole-seedlings nuclei from abovementioned mutants, stained them with DAPI and measured their nuclear area and circularity (Fig.4). Consistently with the other studies (Wang et al., 2013), we observed a reduction in nuclear area and increase in circularity index in *crwn1*, compared to the wildtype control (Col-0). Interestingly, lower average nuclear area was also observed in *pwo1*, albeit with no significant changes in nuclear shape. The analysis on *pwo1 crwn1* phenotype revealed that both genes act in the same pathway to control the nuclear size as the double mutant phenocopies either of the single mutants, rather than displays an additive phenotype (Fig.4B). However, as both, *pwo1* and *crwn1,* show similar nuclear area reduction, we could not determine the phenotypic dominance of one gene over the other in the double knock-out line. Interestingly, we also observed a genetic interaction in nuclear shape phenotype. Namely, as *pwo1 crwn1* shows circularity index average on the similar level as *pwo1*, CRWN1-dependent nuclear shape alteration was suppressed by the lack of functional *PWO1* (Fig.4C). Consistently, we found that the nuclei from *pwo1 crwn1* can be characterized by a broad distribution of circularity index levels, similarly to the *pwo1* single mutant. In contrast, the nuclei from *crwn1* are narrowly distributed and enriched at the high circularity index levels (Fig.4D). Such result suggests that PWO1 is required for CRWN1-mediated control of nuclear shape.

**Fig.4.**
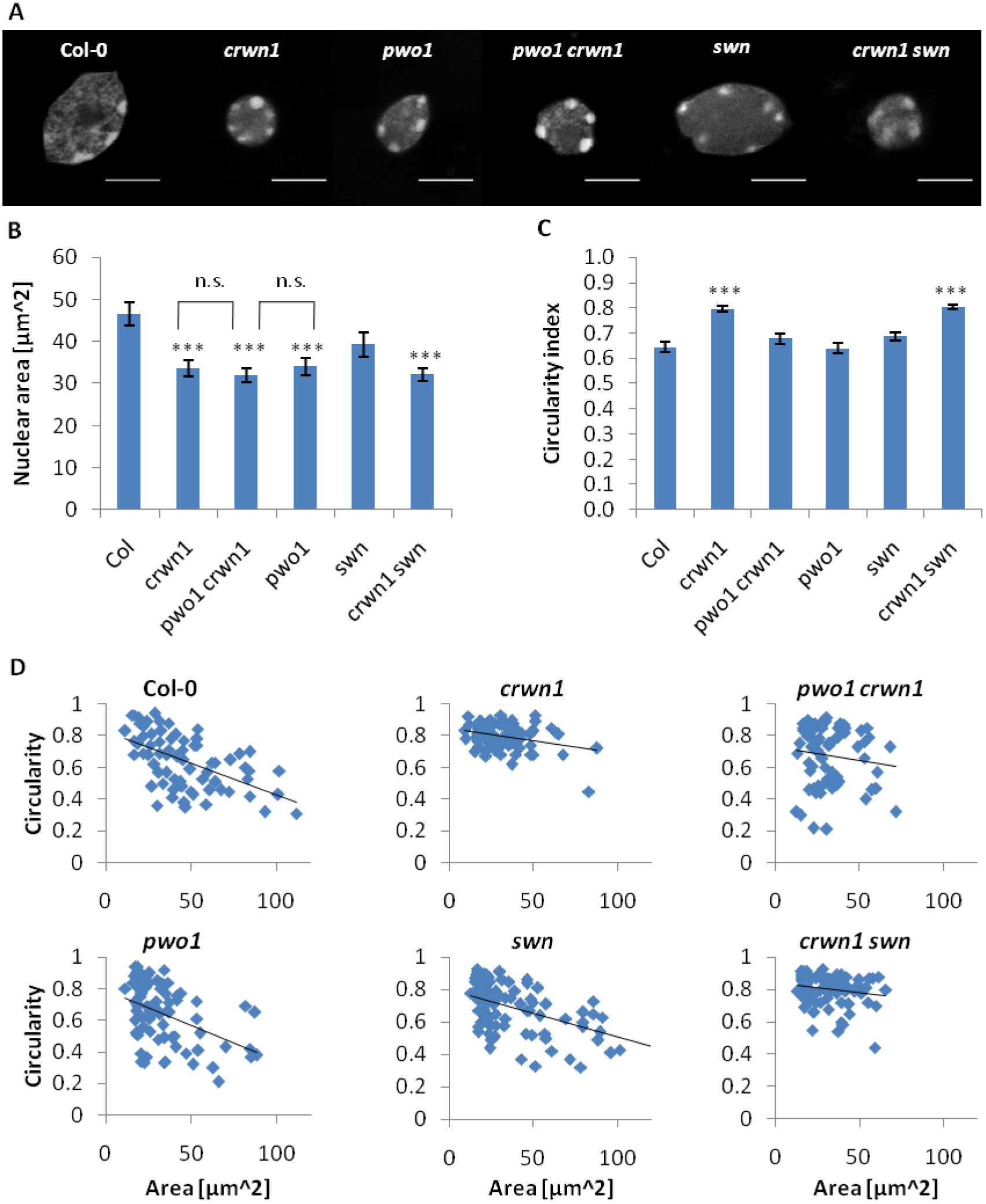
Genetic interaction of *pwo1, crwn1* and *swn*. (A) DAPI-stained nuclei in different genotypes. Scale bar corresponds to 10 μm. Nuclear area (B) and circularity (C) measurements. Calculations were performed on Z plane showing the largest area of particular nucleus in the Z-stack. 72-92 nuclei per genotype were measured. Circularity index was calculated as 4π(area/perimeter^^^2), with the maximal value of 1.0 corresponding to the perfect circle. Significance was calculated in Student’s t-test for comparison of the mutants to the wildtype control (Col). Significance was calculated also for the comparison of *crwn1* to *pwo1 crwn1,* and of *pwo1* to *pwo1 crwn1* (indicated by brackets above respective chart bars). Nonsignificant changes with threshold p = 0.05 are indicated (n.s.), whereas significant changes with p < 0.001 are marked with triple asterisk (***). (D) The relationship between circularity index and nuclear area in nuclei from genotypes used in the study. Blue points represent individual nuclei, black line displays the trendline.

Furthermore, we observed no significant changes in nuclear morphology in *swn*. The double mutant *crw1 swn* shows an increased average circularity and reduced average nuclear area, hence resembling *crwn1*. Consistently, a distribution of individual nuclei in *crwn1 swn* is more similar to the distribution observed in *crwn1*, rather than *swn*. The phenocopy of nuclear morphology changes between *swn crwn1* and *crwn1*, yet lack of such phenotype in *swn*, suggest altogether that the canonical PRC2 doesn’t influence CRWN1-mediated control of nuclear architecture.

Overall, our results suggest that PWO1 and CRWN1 determine nuclear size and shape within the same genetic pathway. In contrast, canonical PRC2 component, SWN, does not influence morphology of the nucleus, irrespective of CRWN1 function.

### PWO1 subnuclear localization

As PWO1 interacts physically and genetically with CRWN1, a nuclear lamina component that localizes to the nuclear periphery, we asked whether both proteins share similar subnuclear localization. In *Arabidopsis*, CRWN1 expressed from its endogenous promoter forms a thin ring underlying the border of the nucleus (Dittmer et al., 2007). If expressed from a strong, 35S promoter in the heterologous system (*Nicotiana benthamiana*), CRWN1 localizes partially to the nuclear periphery, forms ring-like or filamentous structures and induces moderate deformation of NE (Goto et al., 2014).

In order to study PWO1 localization in a heterologous system, we expressed PWO1-GFP fusion protein in *N. benthamina* leaves. In order to prevent formation of over-accumulated protein aggregates, the expression was driven by a beta-estradiol-inducible 35 promoter (i35S). We observed that PWO1 localizes to the nucleoplasm and forms multiple speckles present at the nuclear periphery and the nucleolus (Fig.5, Fig.S3). We detected 17-37 speckles per nucleus (n=13), with a vast majority of them located at the nuclear periphery (Fig.5B).

**Fig.5.**
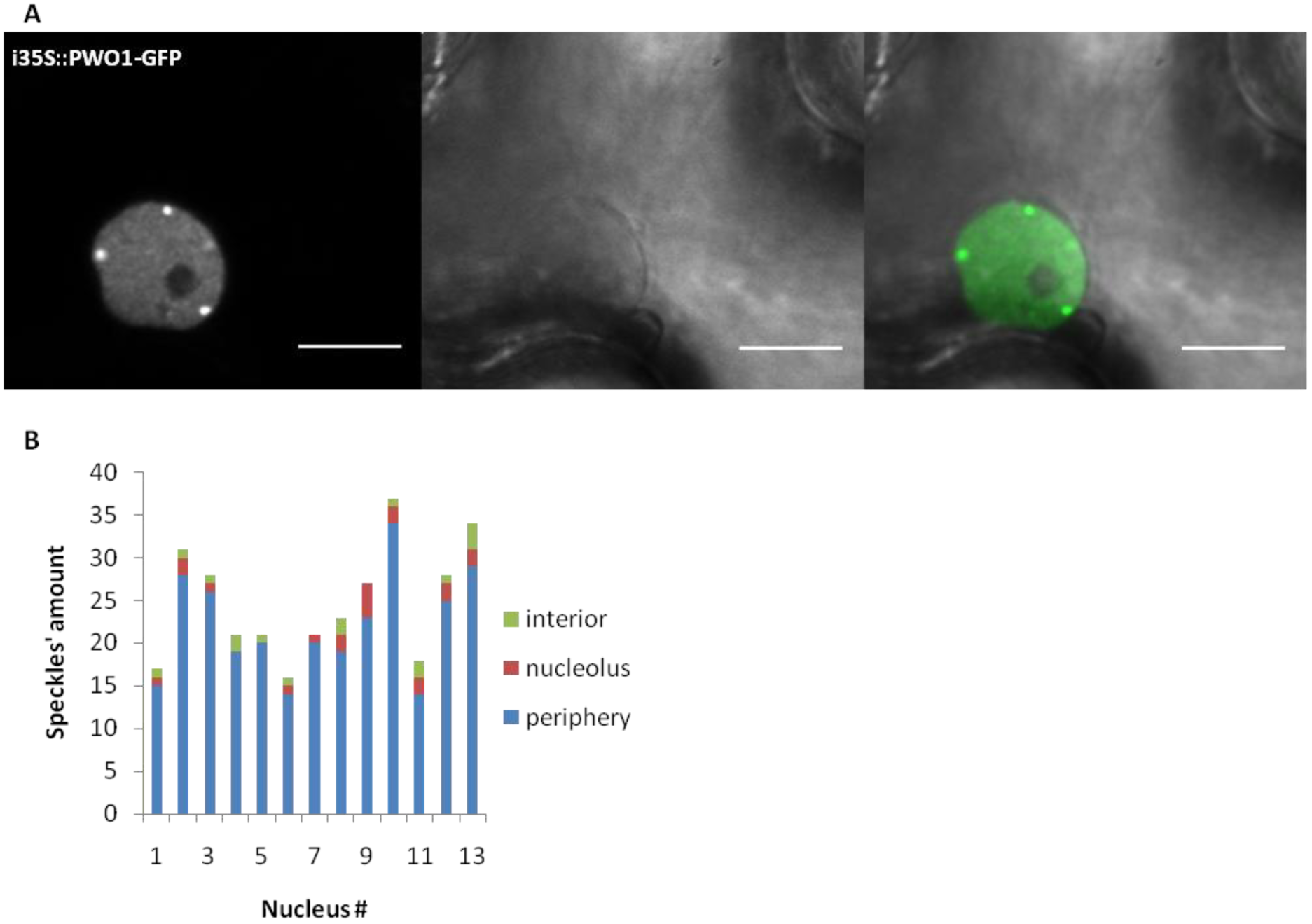
PWO1 localization to speckles in *Nicotiana benthamiana*. Leaves of *N. benthamiana* were infiltrated with beta-estradiol-inducible i35S::PWO-GFP construct. The images of epidermis nuclei were acquired 16-20 hours post induction. Raw images were filtered by Gaussian blur (sigma = 0.75). Scale bar corresponds to 10 μm. (B) The number of speckles and their localization per individual nucleus.

Next, we acquired confocal microscopy images from stable *PWO1::PWO1-GFP Arabidopsis* line. We observed localization to the nucleoplasm and subnuclear speckle-like structures in cortex and epidermis tissues of root meristematic zone (Fig.6A, C). However, due to the small size of meristematic nuclei, it remains unclear whether the speckle-like structures locate preferentially at the nuclear periphery and/or the nucleolus. In contrast, imaging of nuclei from epidermis of root elongation zone and leaves revealed a uniform nucleoplasmic distribution without speckle-like structures’ formation (Fig.S4, Fig.S6A). In summary, we showed that PWO1 does not localize exclusively to the nuclear periphery in *Arabidopsis* or *N. benthamiana.* Instead, it occupies whole nucleoplasmic space and forms partially peripheral subnuclear speckle-like structures in heterologous system and in tissue-specific manner in *Arabidopsis.*

**Fig.6.**
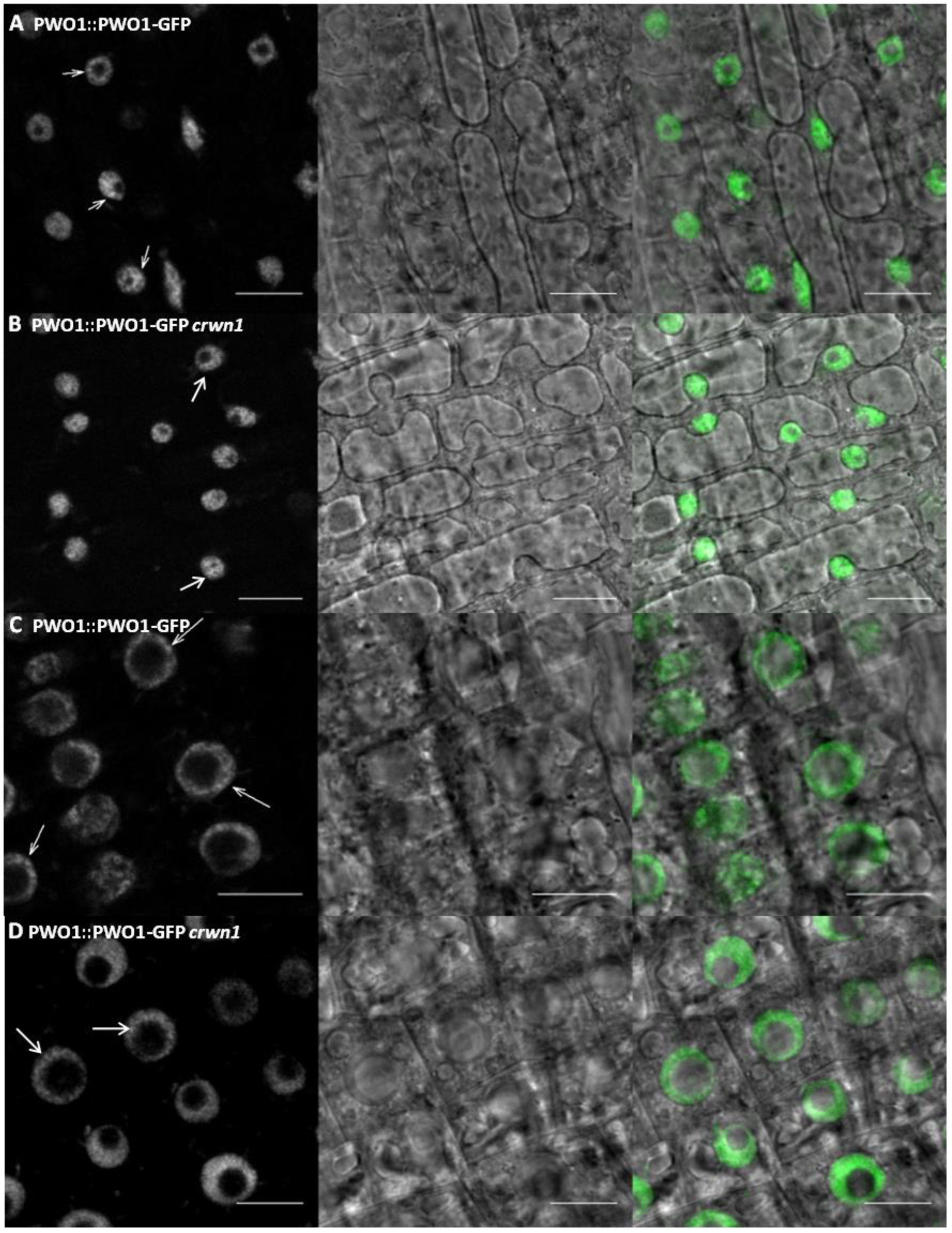
PWO1 localization in *Arabidopsis*. Confocal microscopy was performed on roots from the line: *PWO1::PWO1-GFP* in *crwn1* (B,D) and the control: *PWO1::PWO1-GFP* (A,C). The images represent different root tissues: meristematic zone epidermis (A and B), mersitematic zone cortex (C and D). The images were filtered using gaussian blur (sigma = 1). Scale bar corresponds to 10 μm.

Furthermore, we asked if disruption of *PWO1* or *CRWN1* affects localization of each other. Our confocal microscopy analyses of the lines: *PWO1::PWO1-GFP crwn1* and *CRWN1::CRWN1-GFP pwo1 pwo3* revealed no changes in subnuclear localization comparing to the control lines (Fig.6, Fig.S5, Fig.S6). Therefore, we concluded that the subnuclear localizations of PWO1 and CRWN1 are independent of each other.

### PWO1 and CRWN1 affect expression of a similar set of genes

In order to further understand a connection of PWO1 and CRWN1, we sought to investigate global transcriptomic changes using the RNA-seq method. As a material, we used 2 week-old *Arabidopsis* seedlings from *pwo1* and strong *crwn1 crwn2* mutants. Differential expression analysis (see materials and methods) allowed to identify 179 significantly upregulated‐ and 242 significantly downregulated-differentially expressed genes (DEGs) in *pwo1*, compared to the wildtype control (Col-0). In turn, similar analysis on *crwn1 crwn2* revealed 602 significantly upregulated‐ and 214 significantly downregulated-DEGs. Cross-comparison of the datasets from both genotypes allowed to identify a significant overlap of 71 upregulated‐ and 73 downregulated-DEGs (Fig.7A,B). Such results show that *PWO1* and *CRWN1* regulate partially similar set of target genes.

**Fig.7.**
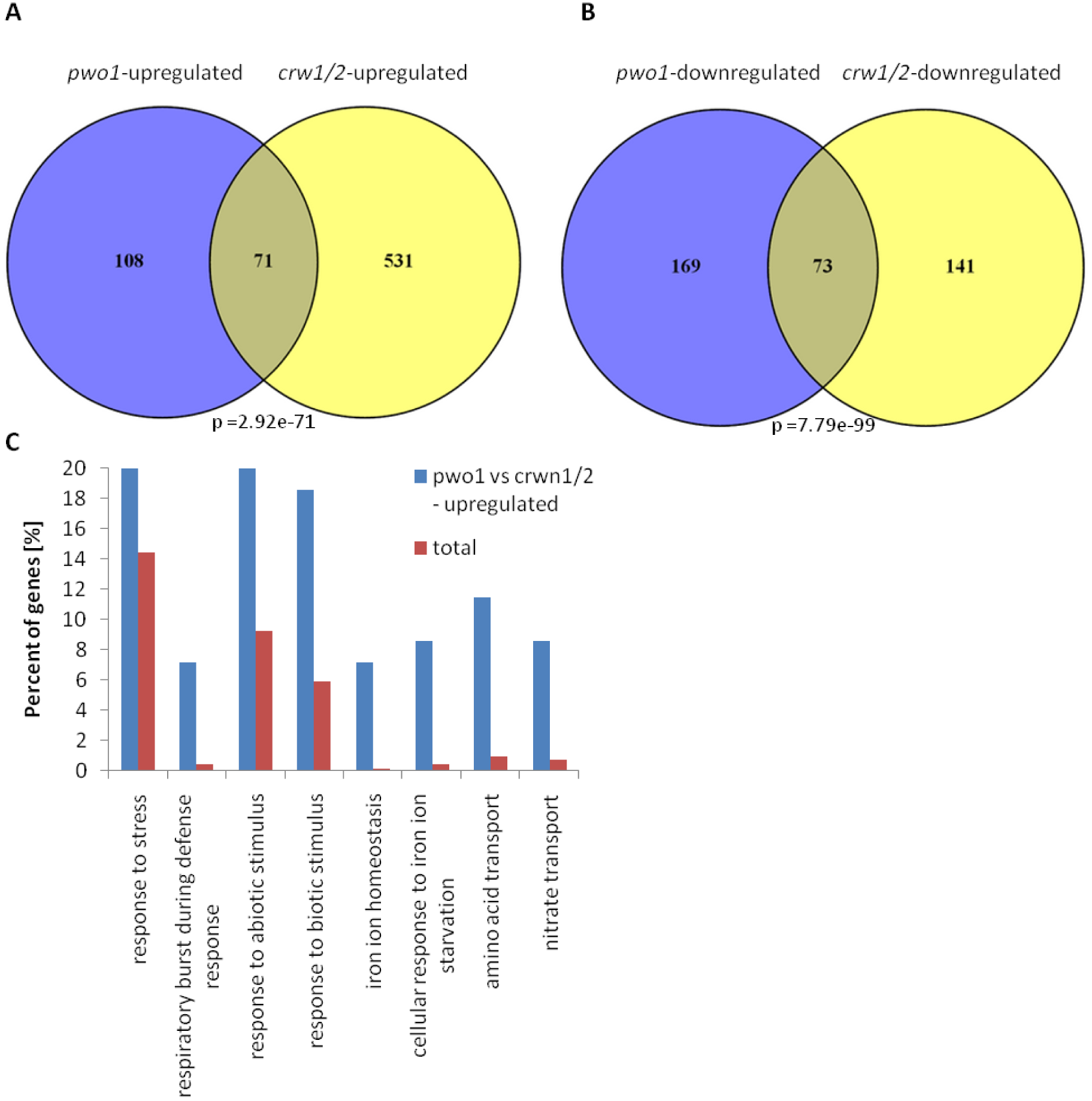
Differentially expressed gene (DEG) analysis – overlap between *pwo1* and *crwn1/2*. (A) Venn diagram comparing upregulated-DEGs in *pwo1* (*pwo1-*upregulated) and *crwn1/2* (*crwn1/2-*upregulated). Significance level of the overlap was calculated in R using hypergeometric test. (B) Venn diagram comparing downregulated-DEGs in *pwo1* (*pwo1-*downregulated) and *crwn1/2* (*crwn1/2-*downregulated). Significance level of the overlap was calculated in R using hypergeometric test. (C) GO term enrichment analysis on commonly upregulated-DEGs in *pwo1* and *crwn1/2* compared to all TAIR10 protein-coding genes (“total”). The percentage corresponds to the number of genes enriched in significantly abundant (FDR<0.005) GO terms in the overlap shown in (A) or the whole protein-coding gene number based on TAIR10 annotation.

Next, we asked whether common DEGs affected by both, *pwo1* and *crwn1/2,*can be organized into functional categories. Our gene ontology analysis (see materials and methods) on the overlap of upregulated DEGs showed significant enrichment of the functions related to: response to stress (biotic and abiotic), iron homeostasis and transport of amino acids/nitrate (Fig.7C). In turn, DEGs from the overlap of downregulated targets are involved specifically in the response to auxin.

Given the PWO1 involvement in PRC2 pathway (Hohenstatt, 2012), we hypothesized that upregulated-DEGs in *pwo1* are covered by H3K27me3 in the wildtype plants, in which a functional PWO1 is present. Therefore, we compared our list of upregulated-DEGs in *pwo1* and published dataset of H3K27me3 targets (Oh et al., 2008). Indeed, we detected a significant overlap of 105 genes present in both lists (Fig.8A,B), showing that PWO1 is required for the repression of a subset of PRC2 targets. As PWO1 associates with CRWN1, we asked whether mutants in CRWN gene family affect expression of H3K27me3 targets as well. Interestingly, the comparison between upregulated-DEGs in *crwn1/2* and H3K27me3-covered genes revealed a significant number of genes shared between two datasets (Fig.8A,B). Finally, cross-comparison of upregulated-DEGs in *pwo1* and *crwn1/2*, and H3K27me3 targets gave 36 genes shared by all three datasets (Fig.8A,B). Their function is related to stress response and amino acid/nitrate transport, but not to iron metabolism (Fig.8C), which implicates a role of PWO1 and CRWN1 in iron homeostasis facilitated in PRC2-independent manner. We noted also a substantial number of H3K27me3 targets which expression is affected only by either, PWO1 or CRWN1 CRWN2. Such a result suggests that PWO1 and CRWN1 mediate repression (directly or indirectly) of PRC2 targets also separately of each other.

**Fig.8.**
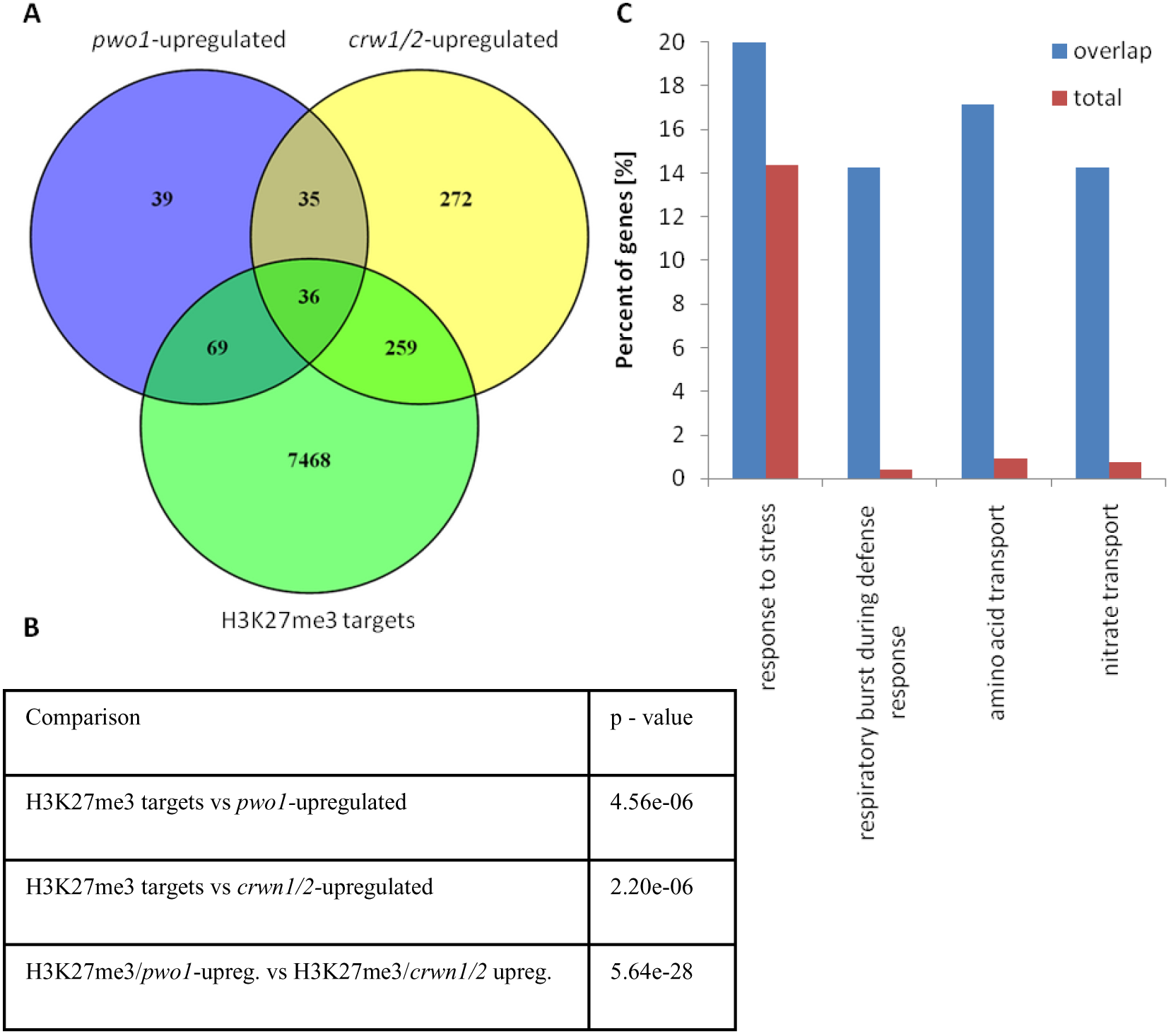
Differentially expressed gene (DEG) analysis – cross-comparison between mutants and H3K27me3 targets. (A) Venn diagram comparing upregulated-DEGs in *pwo1* (*pwo1-*upregulated), in *crwn1/2* (*crwn1/2-*upregulated) and H3K27me3 targets in Col-0 (Oh et al., 2008). (B) Significance level of overlapping gene number shown in (A). P-values were calculated in R by hypergeometric test. (C) GO term enrichment analysis on commonly upregulated DEGs in *pwo1* and *crwn1/2,* and H3K27me3 targets in Col-0, compared to all TAIR10 protein-coding genes (“total”). The percentage corresponds to the number of genes enriched in significantly abundant (FDR<0.0005) GO terms in the common target genes shared by all three datasets in (A) or the whole protein-coding gene number based on TAIR10 annotation.

## Discussion

Here, we demonstrate that the *Arabidopsis* PRC2-associated protein PWO1 interacts physically with numerous nuclear lamina components, including CRWN1 – a coiled-coil analog of lamin proteins in *Arabidopsis*. We report that *PWO1* and *CRWN1* are epistatic in controlling nuclear size and affect expression of a similar set of PRC2 target genes. Moreover, we show that PWO1 partially localizes to the speckle-like structures located at the nuclear periphery.

Our findings suggest a conservation of the association between a repressive chromatin environment and the nuclear periphery across biological kingdoms, including plants. Strikingly, the exact mechanism and linking factors involved in this phenomenon are frequently species-specific, as has been reported for, i.e. *C. elegans*-specific CEC4 (Gonzalez-Sandoval et al., 2015) or Sir4, present in *Saccharomycetacea e*only (Hickman et al., 2011; Taddei et al., 2004). Identification of PWO1, a novel plant-specific bridging factor, adds another evolutionary-distinct element in otherwise widespread machinery linking gene silencing with the nuclear lamina.

The pathways linking repressive chromatin and NL vary not only between the species, but also within the same organism. Many protein effectors associated with NL in fungi or animals regulate only a specific subset of targets and play roles also outside of the nuclear periphery (Harr et al., 2016). Consequently, our results on PWO1 subnuclear localization, transcriptomic analyses in *pwo1* and *crwn1/2* and whole nucleoplasmic distribution of H3K27me3 in plants (Mathieu et al., 2005) collectively suggest a NL‐ and PWO1-dependent repression of only a specific set of PRC2 targets. We speculate that the other, PWO1-independent, pathways bridging NL and PRC2 also exist, as suggested by the PWO1-independent CRWN1/2-mediated repression of PRC2 target genes. Needless to say, an identification of such pathways will be the essential step for the future. The feature of altered nuclear size in mutants of stable NL components (like *crwn1*) and its associated members (like *pwo1*) provides a measurable phenotype for unbiased mutagenic screens (Goto et al., 2014).

Target-specificity of NL-chromatin bridging pathways is demonstrated also by the phenotypes of respective mutants. Lack of bridging factors is frequently represented by a normal viability and only mild aberrations in standard conditions, with a substantial effect seen only at selected developmental stages or environments (Gonzalez-Sandoval et al., 2015; Ozawa et al., 2006; Towbin et al., 2012). Similarly, the *pwo1* mutant shows general phenotype indistinguishable from the wildtype, with obvious changes known so far only for one specific trait only (flowering time) (Hohenstatt, 2012). As disruption of all PWO family members in *pwo1/2/3* displays severe effect (seedling lethality) (Hohenstatt, 2012), we cannot exclude that the mild phenotype in *pwo1* single mutant is caused by the redundancy between PWO family proteins. However, a nuclear area reduction seen already in *pwo1* single mutant and the absence of conserved CRWN1-interacting motif in PWO2 and PWO3 suggest that PWO1 has a NL-connected role separate from the other PWO family members.

Furthermore, our results on PWO1 subnuclear localization showed formation of peripheral and nucleolar speckles in *N. benthamiana* and whole nucleoplasmic speckle-like structures in *A. thaliana*. Formation of distinguishable nucleoplasmic foci is a prominent feature of PRC1 proteins in animals (Pirrotta and Li, 2012) and was reported to be dependent on PRC2 activity (Hernández-Muñoz et al., 2005). Moreover, animal H3K27me3 is found in large aggregates at the nuclear periphery if associated with late-replicating chromatin (Hernández-Muñoz et al., 2005). Similarly to animal models, PRC1 members in *Arabidopsis* were found to localize into multiple speckles as well (Calonje et al., 2008; Libault et al., 2005), albeit without any preference toward nuclear periphery. Moreover, using genomic methods, H3K27me3 was shown to be enriched in the chromatin associated with nucleolus in mammals (Németh et al., 2010) and plants (Pontvianne et al., 2016). It remains to be elucidated whether PWO1 acts together with PRC1 complex, if PRC1 and PWO1 speckles are functionally connected and whether PWO1 has a role in the organization of nucleolar chromatin.

Our transcriptomic analyses, presented together with the nuclear size phenotype, highlight another important point. As reduced nuclear area causes elevated frequency of interchromosomal contacts (Grob et al., 2014), changes in the gene expression in *pwo1* and *crwn1/2* might reflect indirect effects coming from altered spatial organization of the chromatin. Thus, a generation and cross-comparison of transcriptomic data from the other mutants affected in nuclear size should be performed to de-couple gene-specific influence from the indirect effects.

In short, we provide a putative link between nuclear lamina and H3K27me3-mediated gene repression in plants, highlighting an importance of nuclear architecture proteins in chromatin-mediated control of gene expression across biological kingdoms. Deepening our understanding of PWO1 serves as a fascinating opportunity to decipher mechanistic in the association between gene silencing and nuclear periphery.

## Materials & methods

### Plant material

*A. thaliana* seedlings were grown on ½ MS plates in long day conditions (22°C) and transferred to soil, if later stages of development were needed. For the knock-out mutations, following transgenic lines were used: *pwo1-1* (SAIL_342_C09), *crwn-1* (SALK_025347), *crwn1-1 crwn2-1* (SALK_025347, SALK_076653), *swn-7* (SALK_109121). For transient assays, *N. benthamiana* plants were grown in soil in long day conditions (22°C) up to 4^th^ week and subsequently used for infiltration. The sequences of oligonucleotides used for genotyping are as described in: (Hohenstatt, 2012) and (Wang et al., 2013).

### Cloning

Regarding the vectors used for FRET-APB, pMDC7 derivatives containing GFP, mCherry and GFP+mCherry coding sequences were created as described in: (Bleckmann et al., 2010). Next, genomic sequence of *CRWN1* (AT1G67230), coding region of full *PWO1* sequence (AT3G03140) or coding region of PWO1 C-terminal fragment were ligated into pCR8-GW-TOPO using TA Cloning Kit (Thermo Fisher) to create entry vectors. To construct translational fusion plasmids, insert sequences from the entry vectors were introduced into pMDC7 derivatives via LR reaction from Gateway cloning protocol (Thermo Fisher). For yeast-two-hybrid experiments, CRWN1, PWO1 and PWO1 fragments’ coding sequences were used to create entry vectors via TA Cloning and subsequently introduced into pGADT7 and pGBKT7 Y2H vectors (Clontech) using LR reaction (Thermo Fisher).

### CoIP-MS/MS

Nuclear proteins were isolated from 3g of whole seedlings from *PWO1::PWO1-GFP* or Col-0 (negative control) lines grown in LD conditions. 4 biological replicates per genotype were used. Nuclei were extracted without prior fixation of the tissue as described in: (Kaufmann et al., 2010) and the further procedure was performed as described in: (Smaczniak et al., 2012). For immunoprecipitation, μMACS GFP Isolation Kit (Miltenyi Biotec) was used. Peptide spectra wereobtainedon LTQOrbitrapXLmassspectrometer(ThermoScientific).Protein identification was performed by the database search with Andromeda search engine (Cox et al., 2011). Raw data processing was done on MaxQuant v1.1.1.36 software (Cox and Mann, 2008). To distinguish PWO1-interacting proteins from the background, Student’s t-test with FDR adjustment was performed on label-free quantification (LFQ) values.

### Subnuclear localization and FRET-APB

For FRET-APB and examination of subnuclear localization in *Nicotiana benthamiana*, estradiol-inducible pMDC7 vector derivatives were transformed into Agrobacterium tumefasciens (GV3101 PMP90 strain with p19 silencing suppressor plasmid) and grown on YEB medium plates for 2 days. Bacterial lawn was scraped into infiltration medium (5% sucrose, 0,1% Silwet-L77, 450μM acetosyringone) to OD = 0,8 and kept on ice for 1 hour. Bacteria were infiltrated into 4-week old *N. benthamiana* leaves using 1ml-suringes without needles. After 48 hours, an induction was performed by painting abaxial side of the leaves with 20-50μM Beta-estradiol solution in 0,1% Tween-20. Confocal miscroscopy was done 18-24 hours post-induction on LSM780 (Carl Zeiss) or SP8 (Leica). For FRET-APB, GFP was excited at 488 nm with argon laser and mCherry at 561 nm with helium laser. Photobleaching was performed using 561nm laser in 5 frames with 100% power on Leica SP8. FRET efficiency was calculated as described in: (Bleckmann et al., 2010).

Subnuclear localization of candidate proteins in *Arabidopsis* was done by confocal microscopy on seedling of stable transgenic lines using same laser parameters as described above. Images were acquired with 3-4 line averaging.

### Yeast-two-hybrid

Yeast cultures were grown at 28°C on YPD on selective SD media. AH109 strain was used for transformation, following protocol described in Yeast Protocols Handbook (Clontech Laboratories, Inc. Version No. PR973283 21), with both, Gal4-BD and Gal4-AD, constructs added. Transformants were selected on SD medium lacking tryptophan and leucine (SD-LW). Protein interaction was assessed by the transformant growth on selective SD media additionally lacking histidine (SD-LWH).

### Nuclear morphology analysis

0.5g of 2 week-old seedlings were fixed in 10mL freshly prepared ice-cold 4% formaldehyde in PBS buffer for 20 min. under vacuum. The seedlings were washed 3 times for 5 min in PBS buffer. After removal of PBS, the material was chopped on ice with razor blade in 50μL Nuclear Isolation Buffer (NIB: 500mM sucrose, 100mM KCl, 10mM Tris-HCl pH 9.5, 10mM EDTA, 4mM spermidine, 1mM spermine, 0.1%(v/v) Beta-mercaptoethanol). Additional 450 μL NIB was added and cellular solution was filtered through 50 μm cell strainers (#04-0042-2317, Partec). The filtrate was centrifuged at 500g, for 3 min, 4°C. The pellet was resuspended in 40 μL NIB and 2-3 μL was spread on microscopic slide, left to dry and mixed subsequently with 4 μg/mL DAPI solution (#6335, Roth) in Vectashield mounting medium (#H-1000, Vector Laboratories). DAPI-stained nuclei were visualized in Z-stacks done on SP8 confocal microscope (Leica) using 405 nm diode. Acquired Z-stacks were manually thresholded and used for nuclear shape and size measurements in Fiji implementation of ImageJ2 (Schindelin et al., 2012). Nuclear area and circularity index measurements were performed on Z plane showing the largest area of particular nucleus in the Z-stack. Circularity index was calculated as: 4π(area/perimeterˆ2), with maximal value of 1.0 corresponding to the perfect circle. The measurements were done on 72-92 nuclei per genotype.

### Transcriptomics

RNA was extracted from 2-week old seedlings using RNeasy Plant Mini Kit (Qiagen, #74904), according to manufacturer’s manual. RNA was treated with DNase and its quality was assessed on Nanodrop 1000 (Thermo Fisher Scientific), by Qubit RNA BR Assay (Thermo Fisher Scientific, #Q12210) or using RNA 6000 Nano Kit (Agilent, #5067-1511) on 2100 Bioanalyzer (Agilent). First strand cDNA was synthesized using RevertAid First Strand cDNA Synthesis Kit (Thermo Fisher Scientific, #K1622) with oligo-d(T) primers, according to manufacturer’s manual. cDNA was sent for library preparation and sequencing on HiSeq 2000 (Illumina) at BGI (www.genomics.cn). Alternatively, cDNA was used for RT-qPCR with KAPA SYBR FAST Master Mix (Kapa Biosystems, #KK4600) on iQ5 detection system (Biorad). Expression levels were calculated by application of ΔΔCt method. Sequences of oligonucleotides are shown in Table S2. All the transcriptomic analyses were done in 3 or 5 independent biological replicates for RNA-seq or RT-qPCR samples, respectively.

### RNA-seq analysis

2 x90bp paired-end raw sequencing reads were scored by quality, cleaned and trimmed from 5’ and 3’ends in Trimmomatic v0.35 and FastQC v0.10. Clean reads were aligned to TAIR10 reference genome by TopHat v2.0.12 with mate inner distance (-r) = 20bp, segment length = 30bp and minimal intron length (-i) = 20bp, according to: (Trapnell et al., 2009). Alignment files were processed in two different pipelines: 1) Cufflinks (Tuxedo protocol, (Trapnell et al., 2013)) and 2) EdgeR (Robinson et al., 2010). 1) FPKM values from mapped reads were calculated in Cufflinksv2.2.1 with enabled: reference annotation (-g), multi read correction (-u), fragment bias correction (-b) and minimal intron length set to 20 bp. Cufflinks output was used to create common transcript reference file in Cuffmerge v2.2.1.0 and perform DEG analysis in Cuffdiff v2.2.1 with multi read correction and fragment bias correction. 2) Mapped reads were transformed into counts using HTSeq v0.6.1 and used as input for edgeR v3.3. Features with less than 1 count per mln were discarded. For remaining features differential expression was computed with adjusted p-values <0.05. DEG analysis in either of pipelines concerned comparison between wildtype and mutant (*pwo1* or *crwn1/2)* samples. In order to ensure the stringency of the analysis, only those genes that were present in DEG lists from both pipelines were taken for subsequent steps. The top DEGs in *pwo1* were validated by RT-qPCR using 5 independent biological replicates (Fig.S7). The oligonucleotides used for the validation are depicted in Table S2. RNA-seq data (raw reads and a processed file) was deposited in Gene Omnibus under series number GSE106615.

Hierarchical clustering of the replicates (Fig.S8) was done based on the matrix containing “counts per million” values calculated in HTSeq v0.6.1. Features with less than 1 count per mln were discarded. Heatmap with correlation between replicates was obtained in Heatmap3 v1.1.1 using default methods for distance computing and dendrogram re-ordering.

### Secondary bioinformatic analyses

Gene ontology of DEGs was inferred from Singular Enrichment Analysis on AgriGO server (Du et al., 2010). Statistical significance was calculated using Fischer test with Yekutieli adjustment method and the threshold of FDR<0.01 was applied. *Arabidopsis thaliana* TAIR10 genes were used as a reference.

For cross-comparison of DEGs in different mutant backgrounds, Venn diagrams were created in Venny v2.1 software (Oliveros, 2007). Statistical significance of overlapping gene number was calculated in R using hypergeometric test.

## Authors’ contributions

PM, SF, MH, DS designed the research. DS, KK, GA, MH designed CoIP-MS/MS experiment. MH performed CoIP-MS/MS experiment. CS analyzed MS spectral data. PM and MH performed nuclear morphology measurements. PM and SF performed microscopy analyses on PWO1 subnuclear localization in *Nicothiana benthamiana*. PM did FRET and Y2H experiments, genetic interaction studies, subnuclear localization in *Arabidopsis* analyses and transcriptomic experiments. The manuscript was written by PM and revised by DS.

## Acknowledgements

We would like to cordially thank Prof. Dr. Ruediger Simon (HHU Duesseldorf, Germany) for providing FRET vectors and Keygene N.V. company (Wageningen, Netherlands) for the training in RNA-seq analysis. We thank Dr. Julia Kleinmanns, Dorota Komar and Suraj Jamge for critical revision of the manuscript. This work was supported by European Commission 7^th^ FP project EpiTRAITS, Boehringer Ingelheim Foundation and the CRC973.

